# Simulation-based Benchmarking of Ancient Haplotype Inference for Detecting Population Structure

**DOI:** 10.1101/2023.09.28.560049

**Authors:** Jazeps Medina Tretmanis, Flora Jay, María C. Ávila-Arcos, Emilia Huerta-Sanchez

## Abstract

Paleogenomic data has informed us about the movements, growth, and relationships of ancient populations. It has also given us context for medically relevant adaptations that appear in present-day humans due to introgression from other hominids, and it continues to help us characterize the evolutionary history of humans. However, ancient DNA (aDNA) presents several practical challenges as various factors such as deamination, high fragmentation, environmental contamination of aDNA, and low amounts of recoverable endogenous DNA, make aDNA recovery and analysis more difficult than modern DNA. Most studies with aDNA leverage only SNP data, and only a few studies have made inferences on human demographic history based on haplotype data, possibly because haplotype estimation (or phasing) has not yet been systematically evaluated in the context of aDNA. Here, we evaluate how the unique challenges of aDNA can impact phasing quality. We also develop a software tool that simulates aDNA taking into account the features of aDNA as well as the evolutionary history of the population. We measured phasing error as a function of aDNA quality and demographic history, and found that low phasing error is achievable even for very ancient individuals (∼ 400 generations in the past) as long as contamination and read depth are adequate. Our results show that population splits or bottleneck events occurring between the reference and phased populations affect phasing quality, with bottlenecks resulting in the highest average error rates. Finally, we found that using estimated haplotypes, even if not completely accurate, is superior to using the simulated genotype data when reconstructing changes in population structure after population splits between present-day and ancient populations.

**Availability:** All software used for simulation and analysis is available at github.com/Jazpy/Paleogenomic-Datasim

## 1 Introduction

Unlike modern DNA, ancient DNA (aDNA) is subject to several factors that make its analysis more complicated than present-day DNA. Ancient DNA is damaged by the passage of time resulting in deamination and fragmentation[1], which makes mapping ancient reads to modern references challenging. It can also be contaminated by environmental DNA belonging to microorganisms, or modern individuals of the same species[2]. Despite this, technical and analytical advances such as next-generation sequencing (NGS)[3] and determining which substrates preserve DNA the best[4] have facilitated paleogenomics–the analysis of genomic information from ancient remains. Up to now, paleogenomic studies have contributed to 1) the development of evolutionary biology[5][6], 2) the inference of demographic histories [7][8], and 3) research of ancient pathogens [9].

For example, analysis of ancient human genomes from distinct time periods have been used to infer population movements [10], and these reconstructions are important to explain the genetic structure of present-day human populations. Specific examples of such studies include the characterization of migratory events in present-day Great Britain before Anglo-Saxon migrations[11], the effects of Zoroastrian migrations on the populations of Iran and India[12], genomic changes in European populations following transitions between the Stone, Bronze, and Iron ages[4], and evidence of barbarian migrations towards Italy during the 4th and 6th centuries[10].

As the availability and coverage of ancient genomes increases[13], the usage of haplotype data in paleogenomics will become more common. Considering that currently there are no benchmarks of how well phasing aDNA works as a function of contamination, read depth and temporal drift, it is important to understand how phasing behaves when performed on aDNA data to guide studies that leverage statistical phasing[11][12] to infer haplotypes. In general, there are three main strategies for DNA phasing. Pedigree phasing uses kinship[14] and genotype data for multiple related individuals, but it is rare to have multiple related individuals and information about how they were related in ancient data sets. Read-based phasing[15][14] takes advantage of the fact that alleles belonging to the same read will be in phase with each other. However, the high fragmentation of aDNA makes read-based phasing difficult or computationally intractable. Finally, statistical phasing uses haplotype reference panels or cohorts of related individuals to determine the likeliest phasing of an individual, by reconstructing unphased individuals as a mosaic of other haplotypes. We can further split statistical phasing into reference panel phasing or population phasing[14], depending on the availability of reference haplotypes. Statistical phasing may be the only viable strategy for paleogenomic data.

Reference panel phasing makes use of a known haplotype panel, i.e, a set of high quality haplotypes that describe one or more populations. Based on this panel, the haplotypes of a new individual can be estimated by reconstructing them as a “mosaic” of the known haplotypes[16]. Three of the most used reference panels are those from the *1,000 Genomes* project[17], the Haplotype Reference Consortium[18], and the TOPMed project[19]. For example, the *1,000 Genomes* project gathered the haplotypes of a total of 2, 504 modern individuals belonging to 26 different populations. These populations can be divided into five super-categories: Africans, East Asians, European, South Asian, and admixed populations from the Americas [17]. This reference panel has been used in several studies in modern populations[20], as well as in ancient data phasing studies[11][12][4][10]. When no representative reference panels exist, population phasing is another strategy that does not require phased samples. Population phasing attempts to create an *ad hoc* reference panel by continuously updating the possible haplotypes of a cohort given only their genotypes. However, without knowledge of the underlying haplotype structure, population phasing is more computationally expensive and less accurate.

While a few studies have phased aDNA using the *1,000 Genomes* reference panel, the performance of software that implements statistical phasing[21][16] has not been evaluated for use with aDNA. Factors such as contamination, low read-depth, deamination, and the time elapsed since the time of the ancient samples needs to be considered as haplotype frequencies change with time[22][23]. Even in the best-case scenario where the reference panel individuals are direct descendants of the ancient population that is being sampled, the population might have experienced bottleneck and migration events, that together with temporal genetic drift, could decrease the reliability of phased ancient genomes.

In this study, we developed a pipeline to simulate ancient DNA reads to benchmark the accuracy of phasing aDNA. Our simulations account for demographic history, varying levels of contamination, damage, and coverage. We called variants on the simulated data and tested the accuracy of the haplotypes estimated by SHAPEIT[16]. We then measured how well population structure could be reconstructed from these inferred haplotypes. For each demographic scenario, we varied the age of the samples, and the divergence time between the ancient and present-day samples that are used as reference populations. Our results show that increased contamination and lower read depths always lead to elevated phasing error. We found that when the phased individuals belong to the same population as the reference panel individuals, and the samples have a high coverage and little contamination, phasing accuracy is high even with very ancient samples, suggesting that temporal drift is one of the smallest factors affecting phasing accuracy. We found that population splits and bottlenecks have an effect on phasing accuracy. We found that population phasing performs worse than reference panel phasing, and is considerably more computationally expensive. Finally, we used PCA plots on SNP matrices and *ChromoPainter*[24] matrices (which require phased haplotypes and indicate haplotype sharing) to evaluate whether we observed the expected structure. In summary, this work provides useful guidelines for the phasing of ancient individuals, and a tool that can be used for simulating aDNA reads under a user-specified demographic history.

## 2 Methods

### 2.1 Software Pipeline Overview

We developed a pipeline to simulate ancient DNA comprising data simulation, data processing, phasing, and population structure reconstruction (Figure 1, panel A). This software is available as an online GitHub repository[25].

**Figure 1:**
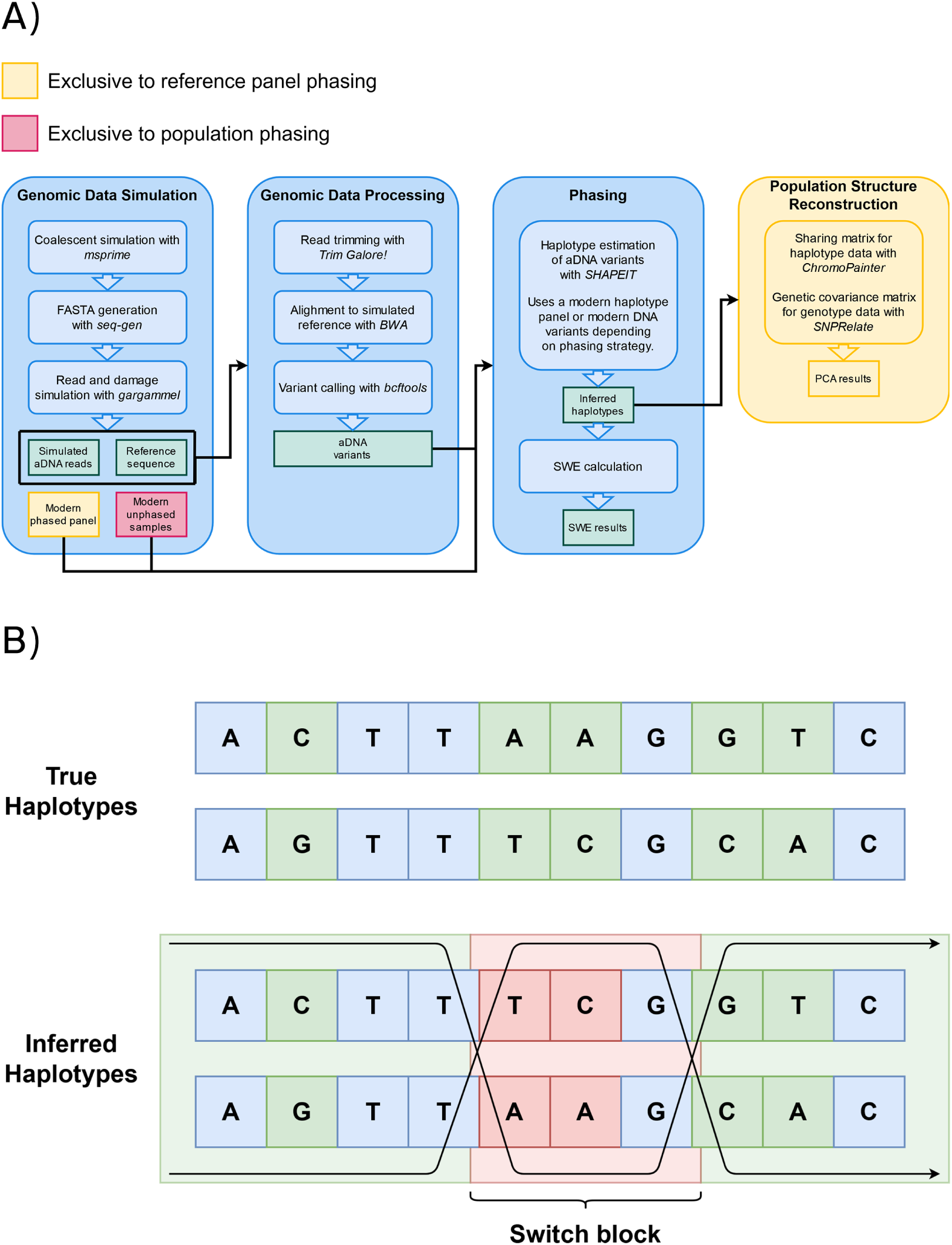
**A)** Data simulation, processing, and analysis workflow. We consider four main stages: data simulation, processing, phasing, and population structure reconstruction. The steps shown inside each stage list the tools required for its execution. Each stage’s execution, input, or output can vary depending on the phasing method chosen (yellow or red). SWE stands for *switch error rate*. **B)** Diagram representation of switch-errors. A switch-error is any “flip” in what the correct maternal and paternal haplotypes should be. Every switch block corresponds to two switch errors, one for each flip. A switch-error rate is calculated as the amount of switch-errors divided by the amount of sites where a switch-error could have occurred. In this example, the phase changes two times across 5 heterozygous sites, resulting in a SWE of 40%.

This pipeline was developed with the intent of being as close as possible to the real workflow of a genomicist working with aDNA, specifically phasing and demographic structure reconstructions using the resulting haplotypes. The pipeline is highly parallelized, and can be easily customized by the user in different ways: the structure, history, and parameters of the simulated samples, the processing of the raw generated sequences, and the application of other methods that aren’t necessarily haplotype phasing.

### 2.2 Genomic data simulation

To generate genomic data, we first use the coalescent simulator, *msprime*[26] with varying demographic models (we consider three models, see Figure 2 and Methods section 2.5). For all simulations, the mutation and recombination rate were set to a value of 2 × 10^*−*8^ per base pair per generation. The length of the sequences was 5 MB. For each simulation, we used the generated coalescent trees as input for seq-gen[27] to generate *FASTA* files for the ancient and present-day individuals. We generated 100 ancient and 502 present-day individuals. Of the 502 present-day individuals, one was used as the reference genome to map reads against and another one was used to introduce contamination into the ancient reads. These two individuals are from exactly the same population as the 500 present-day individuals, but are not part of it. The 500 present-day individuals served as phased reference panel or unphased reference population for reference phasing or population phasing, respectively.

**Figure 2:**
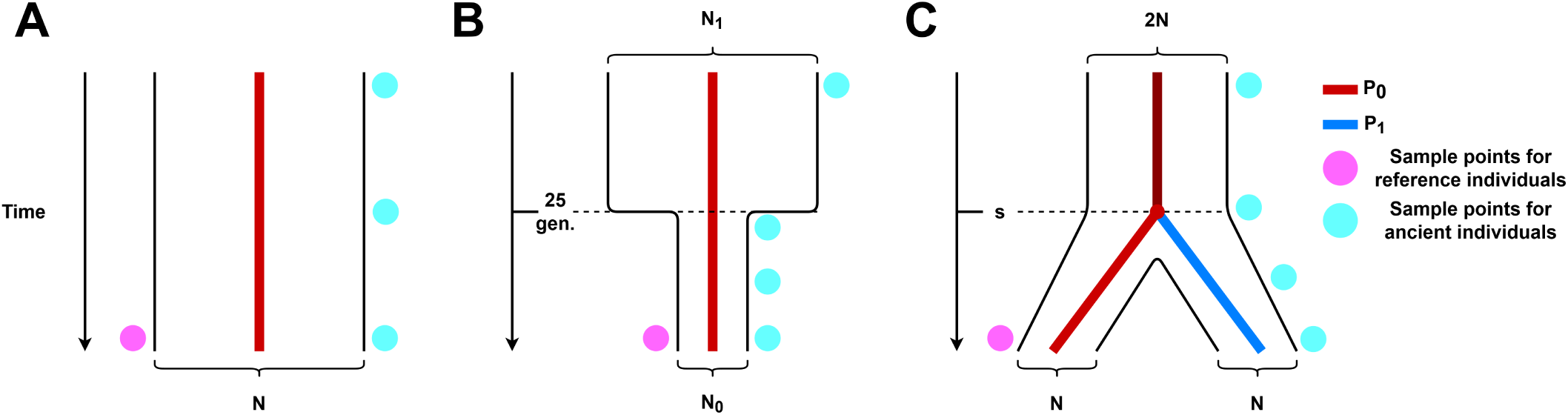
Simulated demographic histories. **A)** No demographic events, we consider a single population *P0* that remains constant in size *N* through time. **B)** Population bottleneck.We consider a single population *P0* that underwent a drastic decrease in population size 25 generations ago. **C)** Population split. We consider two populations, *P0* and *P1*, of equal size *N* that coalesce into a single population *P0* at time *s*, the size of *P0* previous to this time of coalescence is a constant *2N*.

To generate ancient reads, we fed the 200 ancient simulated chromosomes into *gargammel*[28]. This tool introduces damage and fragmentation based on empirical distributions, while also simulating the desired read depth, contamination, and sequencing errors. Table 1 shows the range for each of these parameters that result in 72 different parameter combinations. Simulated average read depths ranged from 1× to 10×, and contamination ranged from 0% to 10%. The values for coverage are illustrative of the data commonly used in aDNA studies. Depth of coverage between 1× and 10× can represent most of the data used in the previously mentioned studies[11][12][4][10], and is also representative of the actual coverage of most ancient genomes sequenced[3][29]. While most of these studies did not contain samples with modern contamination higher than 2%, we also considered values of 5% and 10% to better understand how contamination affects haplotype estimation. The damage profile of different ancient samples depends greatly on environmental factors like temperature and humidity, so it is difficult to confidently create damage profiles that correspond to different sample ages. Because of this, all samples we simulated were damaged with the same default damage matrix available in *gargammel*.

**Table 1:** Parameters for different simulated sample histories.

| Demographic history | No events | Bottleneck | Population split |
| --- | --- | --- | --- |
| Population size ($N$) | 10,000 | 10,000 | 10,000 |
| Post-bottleneck population size ($N_1$) | $N/A$ | 1,000 | $N/A$ |
| Time of event (generations ago) | $N/A$ | 25 | 50, 100, 200 |

Simulated sequencing reads were paired-end trimmed using *Trim Galore!*[30] with default parameters. We then aligned the simulated reads to the simulated reference using *BWA mem*[31] with default parameters. After aligning all reads, we called variants using *bcftools mpileup*[32] and created a VCF file for each individual. Variants were filtered to have a minimum genotype quality score of 20.

### 2.3 Phasing of simulated data

Previous studies that perform statistical aDNA phasing approach this task in two ways. For example, Gnecchi-Ruscone et al. (2022)[33], used a haplotype reference panel built from modern samples to phase aDNA. On the other hand, some studies use population phasing[34][12] by grouping the genotypes of the ancient individuals of interest with a larger number of unphased modern individuals.

We emulate these two types of phasing approaches with *SHAPEIT*. For reference panel phasing we create a phased reference panel directly from the simulated sequence data of the present-day individuals (1000 chromosomes). In population phasing, the phases of present-day samples are ignored and inferred jointly with the phases of ancient individuals. It is much more computationally expensive since all individuals in the merged VCF must be phased.

In both cases, all ancient individuals were phased independently of each other, in other words, the phasing algorithm only had information for the reference individuals plus one specific ancient individual.

### 2.4 Phasing accuracy and switch-error rate

To measure phasing accuracy, we use the inferred haplotypes obtained from *SHAPEIT*, and compare them to the actual haplotypes obtained from the coalescent simulation. We measure the switch error rate (SWE) for each individual, which represents the amount of errors in the estimated haplotypes as a percentage (see Figure 1, panel B), specifically, the amount of sites where a phase change occurs divided by the number of sites where it could occur (heterozygous sites kept after filtering). We obtain a distribution of SWE for each different quality parameter combination.

### 2.5 Demographic events

We tested the accuracy of phasing under three simulated demographic scenarios (Figure 2): a single population of constant size through time, a single population that undergoes a bottleneck event 25 generations in the past, and the case where the modern and ancient individuals belong to two different populations that split at some point in the past *s*. All simulated parameters are listed in Table 1.

The parameters (time and effective population size change) of the bottleneck event were chosen to resemble the magnitude of the population collapse that occurred in some Native American populations due to European colonization 500 years ago[35]. For the population split times, we selected a range of 50, 100 and 200 generations ago which is roughly 1000, 2500 and 5000 years ago. We note that for the population split (Figure 2, panel C), the samples of the reference population are not always descendants of the ancient individuals. We did this to test the effect of using a reference population that diverged at some time in the past from the ancient individuals sampled. For each of these demographic histories, we have 72 possible combinations of sample quality parameters detailed in Table 2, which results in 216 different simulation scenarios. We make 100 replicas under each simulation scenario to have a distribution of our results; leading to a grand total of 72 × 3 × 100 × 2 = 43, 200 phased genomes.

**Table 2:** Tested values for each simulation quality parameter.

| Quality parameter | Simulated values |
| --- | --- |
| Age (generations) | 0 (present-day), 25, 50, 100, 200, 400 |
| Depth | 1×, 5×, 10× |
| Contamination | 0%, 2%, 5%, 10% |

### 2.6 Reconstruction of population splits with phased and unphased data

We employed both Principal Component Analysis (PCA) and *ChromoPainter* for this analysis. We generated longer (20 MB) sequences, since *ChromoPainter* expects sequences that are closer in length to a full chromosome. We only use the haplotypes inferred through reference panel phasing, as they had lower SWE and the running time for population phasing was prohibitively long for the number of replicates needed. We only considered the simulations with an average read depth of 10× and 5×, since 1× data had too few variants left after applying quality filters.

We tested the demographic scenario of a population split 200 generations in the past. PCA was applied to 4 different kinds of data: true genotype data (unphased SNPs), true haplotype data, genotype data called from simulated read data and the corresponding inferred haplotypes. When using genotype data, the genotype covariance matrix was built directly from the unphased VCF file using the *R* package, *SNPRelate*[36]. When using haplotype data, PCA was applied to matrices built with *ChromoPainter*[23] that indicate similarity between samples through lengths of IBS tracts.The first and second PCs were plotted (Figure 5) and we measured how well the modern and ancient populations clustered by computing the *silhouette coefficient*. This metric evaluates clusterings using information inherent to the dataset, so that clustering can be compared across different simulated datasets[37]. Measuring cluster distinction is important, since distance metrics in component space can be thought of as proxies for population split times[38].

## 3 Results

### 3.1 Performance of Population and Reference Phasing

We compared the performance of population versus reference phasing. The lack of phased reference individuals for population phasing decreases the available information as we only have genotype information for the 500 modern individuals, this in turn increases the SWE. We simulated individuals under a bottleneck event 25 generations in the past (Figure 2, panel B), and applied population and reference panel phasing to the resulting data (Figure 3).

**Figure 3:**
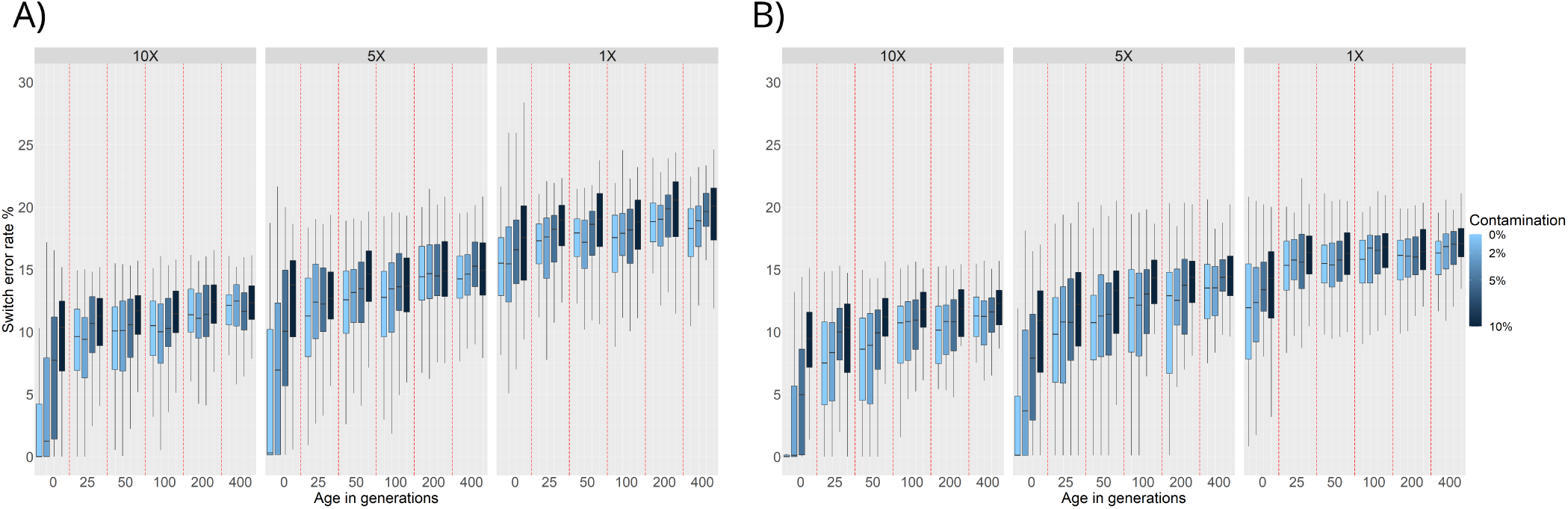
SWE distributions for phasing of ancient individuals simulated under a bottleneck model 25 generations in the past: **A)** Population phasing. **B)** Reference panel phasing. SWE (y-axis) is presented in 3 facets corresponding to 10×, 5×, and 1× average read depth. Within these facets are the SWE distributions for 100 simulated individuals for each combination of age (x-axis) and contamination level (shades of blue).

We find that, as expected, increasing the age and contamination or decreasing the coverage of the simulated samples results in higher SWE. Samples from before the bottleneck event (25 or more generations of age) show a high SWE (*∼* 10%), this can be attributed to the lack of representation of pre-bottleneck haplotypes in the post-bottleneck reference population. These trends are the same for both population and reference panel phasing, however, we can see an overall increase of SWE in the population phasing results compared to the reference panel phasing results.

Another factor to consider is running time as population phasing is more computationally expensive. *SHAPEITv2*’s algorithm has a time complexity of *O(MJ*)[16], where *M* is the number of SNPs to phase, and *J* is the number of haplotypes being conditioned on to build the likeliest phase reconstruction. Phasing a single sample with a reference panel means that *J* = 1, while phasing 501 individuals via population phasing means that *J* = 501. While execution time for phasing all data in one of our simulations using reference panels might take a couple of hours, population phasing on the same data could take upwards of 5 days depending on hardware.

Because of the increase in overall SWE when using population phasing, plus the computational complexity factors, we decided to focus on reference panel phasing results for the rest of the results.

### 3.2 Phasing accuracy as a function of demographic history

Using reference panel phasing, we next consider the effects of demographic history on phasing accuracy. Under a constant population size, we observe an increase in SWE from *∼* 1.0% — when the simulated individuals are from the present (0 generations), have high coverage and no contamination — to *∼* 10.0% when increasing the age to 400 generations (Figure 4, panel A). Increasing the amount of contamination for the individuals with 0 generations of age and high coverage results in higher SWE (*∼* 8.0%). Consistently, decreasing the average read depth from 10× to 5× results in higher SWE (*∼* 1.0% to *∼* 6.0% for modern uncontaminated individuals across the board, and the effects of higher ages and contamination rates are preserved (Figure 4, panel A). Finally, the results for 1× read depth show an increase in SWE across all ages and contamination rates. Within these 1× simulations, the age and contamination show little impact. This likely reflects that individuals sequenced at 1× have a small number of variants and some individuals are excluded because no variants pass all quality filters applied.

**Figure 4:**
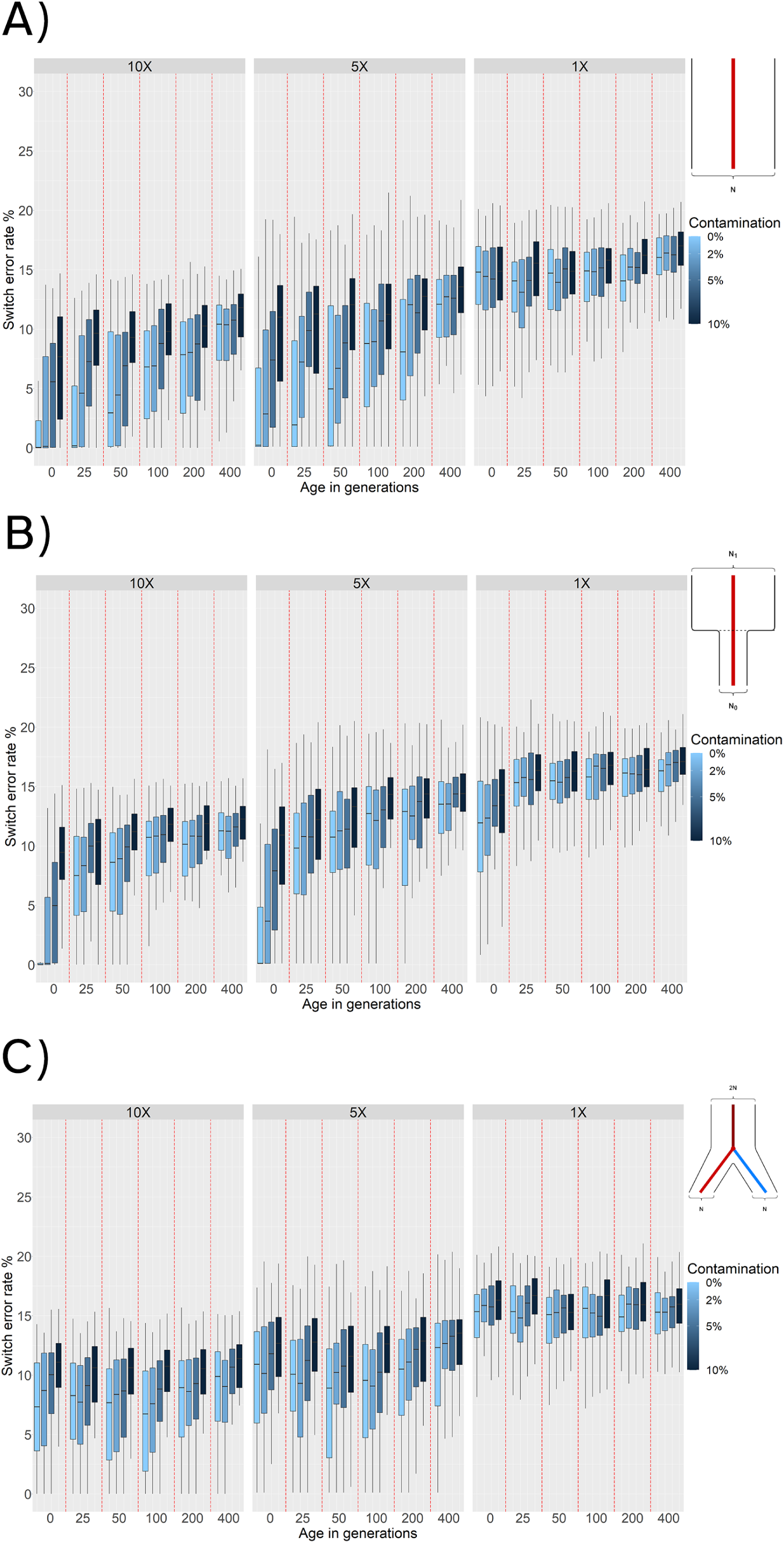
SWE distributions for reference panel phasing with: **A)** Constant population scenario. **B)** Bottleneck event 25 generations in the past. **C)** Population split 100 generations in the past. SWE (y-axis) is presented in 3 facets corresponding to 10×, 5×, and 1× average read depth. Within these facets are the SWE distributions for 100 simulated individuals for each combination of age (x-axis) and contamination level (shades of blue).

To test the effects of a bottleneck, we simulated a population that experienced a 90% reduction in effective population size (*N*_e_ = 10, 000 *→* 1, 000) 25 generations in the past (Figure 2, panel B). We find that phasing quality increases for ancient individuals that are more recent than the time of the bottleneck (0 generations of age, SWE *∼* 0.3%). However, phasing quality for sampled ancient individuals older than the bottleneck event (25 generations or more) decreases. This is expected as individuals older than the time of the bottleneck belong to a population with a much higher diversity that was lost and is not captured by the modern reference individuals. Individuals with an average read depth of 1× exhibit a much higher SWE compared to results with other demographic histories (Figures 4, panels A and C), and we observe that contamination does not have a strong effect for sampled individuals that are older than the time of the bottleneck.

When we evaluate the behavior of phasing individuals of a population undergoing a split 100 generations ago from the population used as the haplotype reference panel (Figure 2, panel C), we observe that ancient individuals sampled around the time of the split (i.e. 50 to 100 generations in the past) exhibit the lowest SWE for 10× and 5× coverage values. This is expected, as samples that are more recent than the time of the split do not belong to the same population of the reference individuals (Figure 2, panel C). Therefore, both more recent and more ancient samples have higher genetic drift from the reference population than the individuals at the time of split. For sequencing coverage of 1×, the phasing quality is lower than with 5× or 10×. In the case of 1× coverage, the effect of age and contamination are negligible suggesting that coverage is the biggest factor for phasing accuracy.

### 3.3 Visualizing population structure

We further tested if population structure could be accurately recovered from inferred haplotypes, and how much of an impact would haplotype estimation error have on these reconstructions. To do this, we simulated samples under a population split model with a population split 200 generations ago (Figure 2, panel C), and sequence length of 20 MB. We sampled 100 ancient individuals 25 generations ago and 500 present-day reference individuals. We note that the ancient samples and reference samples do not belong to the same population in this scenario (Figure 2, panel C). To test if population structure was visually recoverable, we plotted the first and second Principal Components (PCs) obtained by running PCA on four different kinds of data resulting from this simulated demographic history, note that in this context, *true* refers to the exact outputs of the coalescent simulations: (1) the true genotypes, (2) the true haplotypes, (3) genotype data called from the sequencing reads generated with damage, contamination (0-10%) and two read depths (5×, 10×) and (4) the inferred haplotypes using reference panel phasing on the called genotypes from (3). We applied PCA to the genotype matrices (1 and 3) or to the matrices obtained from *ChromoPainter*[23] based on haplotype data (2 and 4; see Methods section 2.6).

Using either the true genotype data (Figure 5, panel A), or the true haplotype data (Figure 5, panel B), PCA reveals distinct clusters for modern reference and ancient samples, which is expected given that they belong to two distinct populations. The clusters are more distinct when recovered from the true haplotype data.

**Figure 5:**
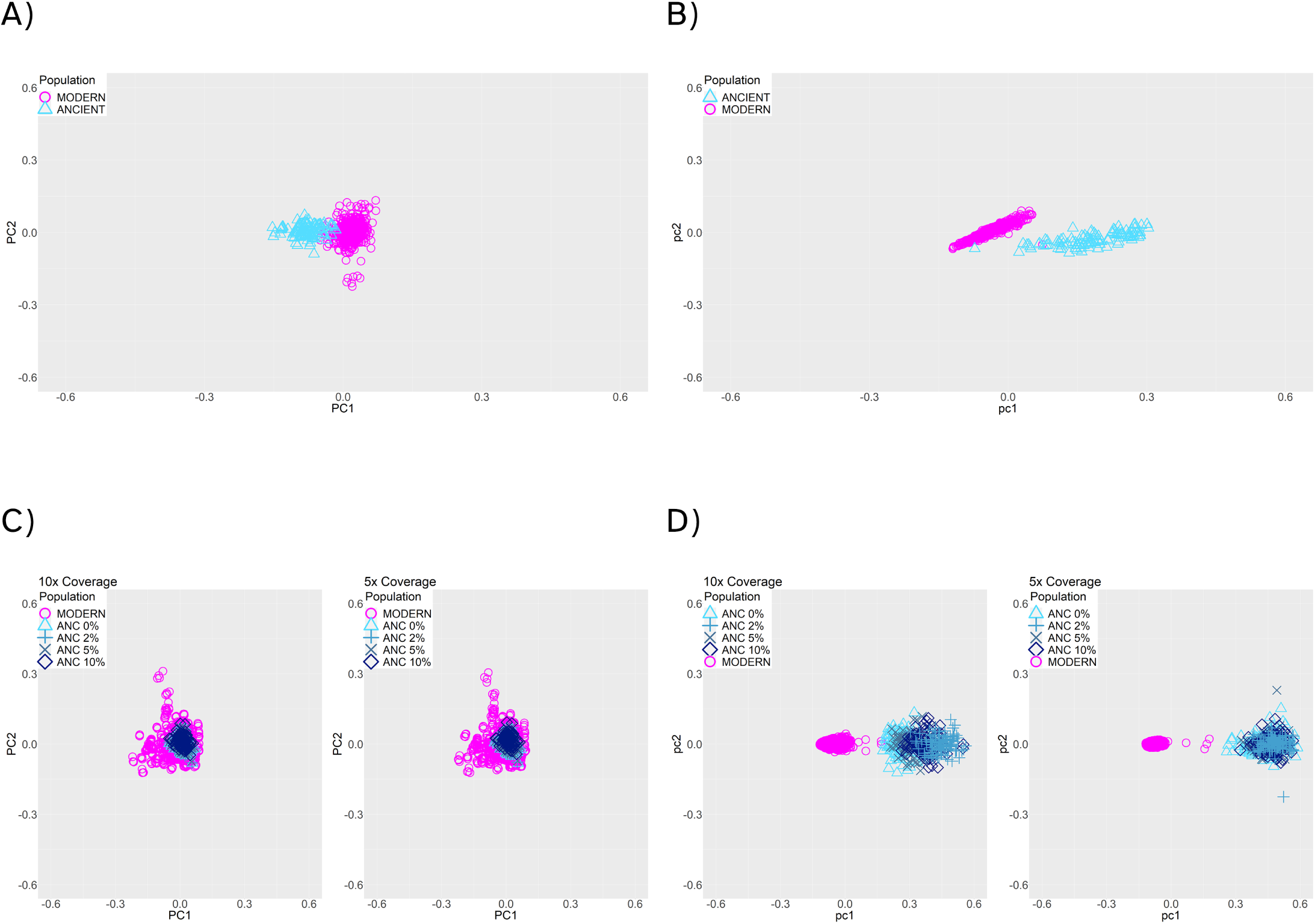
PCA of modern (pink) and 25-generation-old individuals (blue), with a population split 200 generations in the past. **(A)** True genotype matrix (scenario 1). **(B)** True haplotype data (scenario 2). **(C)** Called genotype data. (scenario 3) **(D)** Estimated haplotypes from the called genotype data (scenario 4). Panels **C** and **D** show results with 10× (left) and 5× (right) read depth, and 0% to 10% contamination rates (blue symbols)

When using genotype data after applying damage, contamination, and missingness to the simulated data (scenario 3), the first two PCs no longer recover population structure (Figure 5, panel C). In contrast, the haplotypes recovered after reference panel phasing render a visually recognizable separation between the clusters (Figure 5, panel D), suggesting that using haplotype data provides better resolution. From this, we conclude that the current phasing procedure enables the recovery of population structure from ancient haplotype data, despite the noise present in aDNA.

Silhouette coefficients offer a way of measuring and comparing the clustering performance for these four scenarios, independently of the fact that each PCA was done on a different set of data. These coefficients can range from *−*1 to 1, with values closer to 1 indicating better clustering performance. We found that silhouette coefficients for the PCAs on genotype data (scenario 1 coefficient: 0.499, scenario 3 coefficient: *−*0.131) were substantially lower than those for the PCAs on haplotype data (scenario 2 coefficient: 0.729, scenario 4 coefficient: 0.920). This shows that using haplotype data leads to better clustering performance, and using genotype data after simulating quality problems results in strongly overlapping clusters.

## 4 Discussion

Using haplotype data can be powerful to infer demographic history. While it is common to use haplotype data to infer the demographic history of a population using present-day genomes, only a few studies have phased ancient genomes. In this study, we have developed a pipeline to simulate ancient read sequencing data to benchmark phasing of aDNA as a function of different parameters. Specifically, we show how phasing quality changed as we varied coverage, contamination, temporal drift and population split times. We also examined how these parameters affected levels of observable population structure as captured by PCA and *ChromoPainter*.

We first benchmarked the accuracy of population phasing and reference panel phasing. As our results show that reference-panel phasing is more accurate (using measures of switch error rate, SWE) and faster than population phasing (Results section 3.1), most of the benchmarking analyses performed in this study consider reference panel phasing. When we measured phasing accuracy as a function of coverage, we find that decreasing the average read depth leads to higher SWE, since having less SNPs available for the statistical phasing algorithm reduces the certainty at which phase can be inferred (see Figure 4, panel B), in general, sample coverage has the strongest effect on phasing accuracy. Also, as contamination increases, the phasing quality decreases regardless of whether we used reference panel phasing or population phasing(Figures 3 and 4). This makes sense as introducing new variants via contamination will affect the probability distribution of haplotypes estimated by SHAPEIT, and contamination levels as high as 10% introduce more false variants than any other kinds of damage.

To assess the effects of temporal drift, we sampled ancient genomes at different times in the past. Increasing the age of the simulated ancient individuals directly increases the SWE in the phased haplotypes. This occurs when we simulate either a constant population size or when we simulate bottlenecks (Figure 4, panels A and B). We find that low phasing error rates can be obtained from very ancient individuals if we have good quality samples (Average read depth over 5× and contamination below 5%), and reference panels that are representative of the ancient individual. This is most apparent when no demographic changes through time are simulated, thus ensuring more continuity between the reference and phased populations (Figure 4, panel A).

We find that population bottlenecks increase SWE, especially when the sampling time of the ancient individuals is equal to or greater than the time of the bottleneck. This is probably happening because bottlenecks result in a loss of genetic and haplotype variation. Therefore, haplotype inference for ancient individuals older than the time of bottleneck will always result in a low phasing quality (mean SWE of at least 7.5%), independently of other sample parameters.

Under demographic models with population splits, the behavior of phasing error is different. The lowest error rates (Figure 4, panel C) occur when the sampling time of the ancient individuals is closest to the time of the population split. This implies that older samples do not necessarily lead to worse phasing accuracy, but rather genetic distance from individuals in the reference populations. In other words, when we simulated population splits, we found that proximity to the reference population was the most important factor in terms of sample age. This is expected for two reasons: individuals with a more recent age than the time of the split belong to a population that is increasingly divergent from the reference population. Conversely, individuals that are older than the time of split belong to the ancestral population of the reference individuals, but increasing the age further results in more temporal drift that leads to a higher SWE (Figure 4).

We also assess the implications of phasing ancient individuals in the context of population structure. We find that we can recapitulate the population structure with the inferred haplotypes, and that it is better than using only genotype data (Figure 5, panel D). Parameters such as contamination and coverage slightly affect clustering, but even in those cases we recover population structure (Figure 5).

In this work, we provide the first study (to our knowledge) that benchmarks phasing in aDNA while accounting for various features of the data such as age, contamination, coverage, and demographic history. Although we simulated only a subset of possible data quality parameters and demographic scenarios, these results are a good starting point for guiding future studies that necessitate aDNA phasing. While pieces of the pipeline already exist (e.g. sequence[27] and read[28] simulation), here we provide an easy to install and open source software that streamlines all steps from simulation under a demographic model to visualization which will be helpful for others to evaluate how phasing quality might be affected by characteristics specific to other systems or populations.

## Funding

This work was supported by the PAPIIT program offered by UNAM-DGAPA with filing number IA203821, by grant CN 17-12 of the UC MEXUS-Conacyt program, by the Alfred P. Sloan Award, by a Young Investigator’s grant from the Human Frontier Science Program, and by NIH grant R35GM128946.

## Notes

### Competing Interest Statement

The authors have declared no competing interest.

### Summary of Updates

Corresponding author list.

